# Methionine-triggered growth arrest reveals activation of Gcn2 by methionine transporter endocytosis

**DOI:** 10.1101/2025.05.12.653625

**Authors:** Nathaniel L. Hepowit, Hayley L. Singkhek, Derek J. Johnson, Jason A. MacGurn

**Affiliations:** Department of Cell and Developmental Biology, Vanderbilt University, Nashville, United States

## Abstract

Cell growth checkpoints require coordination between multiple sensing and signaling systems to ensure that cells only proceed with growth and division when conditions are favorable and adequate resources are available. This coordination between nutrient sensing and growth signaling is fundamental to understanding how nutrient supply regulates the cellular metabolic economy. Much of our current understanding is driven by studies that examine the cellular response to nutrient deprivation. For example, TORC1 activity promotes cell growth when amino acids are available, but amino acid deprivation decreases TORC1 activity resulting in activation of catabolic activities. In this study, we examine how cells respond to stimulation with excess amino acids. We report that stimulation with excess Ile, Phe and Met slows cell growth and triggers a G1 cell cycle arrest. Similar to a starvation response, surplus Ile, Phe and Met induce autophagy and trigger decreased TORC1 activity. In the case of stimulation with excess Met, the Gcn2 pathway is required for growth arrest, autophagy induction, and TORC1 dampening. Unexpectedly, Gcn2 is activated by stimulation with excess Met, and this activation requires endocytosis of the methionine transporter Mup1. These results indicate that endocytosis of an amino acid transporter is required to activate the Gcn2 pathway, providing an example for how nutrient transporter trafficking may function as a sensor contributing to cell growth control.

## Introduction

Regulation of metabolism and cell growth decisions requires precise monitoring of nutrient availability and complex coordination between multiple biological systems. For example, the START checkpoint in *Saccharomyces cerevisiae*, which governs the G1/S transition, integrates information on availability of nutrients such as glucose and amino acids to ensure the cell does not commit to division without sufficient resources. The equivalent G1 checkpoint in mammalian cells, known as the restriction point, integrates information on nutrient availability as well as extracellular cues such as growth factors. Precisely how these complex checkpoints process multiple inputs into a coordinated responses remains incompletely understood.

The best understood examples of how cells coordinate nutrient sensing and cell growth decisions involve systems for sensing the availability of glucose. When glucose is available in the environment, yeast prioritize aerobic fermentation, but when glucose in the environment is exhausted cells undergo a diauxic shift to a slower growth program based on aerobic respiration. Several signaling mechanisms contribute to the diauxic shift. At the plasma membrane, Gpr1, a G-protein coupled receptor, responds to glucose by stimulating intracellular cAMP levels, which in turn promotes the activity of PKA (Rødkaer and Faergeman, 2014). Snf3, which also localizes to the plasma membrane, responds to low levels of glucose and signal to alter expression of glucose transporters at the cell surface (Kim et al., 2013). The Snf1 kinase complex, which is homologous to the AMP-activated protein kinase in mammalian cells, responds to declining glucose levels by coordinating the metabolic shift towards respiration. Together, these pathways coordinate the cellular response as glucose becomes limiting in the environment.

Cell growth decisions also require input from systems that sense and respond to the availability of nitrogen sources such amino acids and nucleic acids. One well-studied example involves the TORC1 kinase complex, which signals to generally promote macromolecular biosynthesis (e.g., translation) and to inhibit catabolic activities (e.g., autophagy). Regulation of TORC1 activity has been studied extensively and involves multiple upstream protein complexes, some of which sense and respond to availability of amino acids (either directly or indirectly) (Cui et al., 2023). Another key nitrogen starvation response mechanism involves phosphorylation of phosphorylation of eIF2α, which generally inhibits global translation initiation while simultaneously derepressing translation of Gcn4, a transcription factor that activates a starvation response which includes expression of genes involved in amino acid biosynthesis. This is referred to as the integrated stress response, because several stress-responsive kinases phosphorylate eIF2α in mammalian cells. In yeast, the only kinase known to phosphorylate eIF2α is Gcn2. Gcn2 is conserved across eukaryotic evolution and is activated by uncharged tRNA which accumulate during amino acid deprivation (Dong et al., 2000). Activation of Gcn2 involves the regulatory Gcn1-Gcn20 complex, although many details regarding the activation of Gcn2 remain unclear.

While the effects of nutrient deprivation have been studied for decades, less is known about how cells sense and respond to nutrient surplus. Here, we report our findings that certain amino acids, when provided in excess, trigger a response which mimics starvation. Specifically, we find that excess Ile, Phe and Met triggers a slow growth program associated with G1 cell cycle arrest and the activation of autophagy. Characterization of the autophagic response revealed differences in kinetics and cargo specificity of the response depending on the amino acid stimulus. Stimulation with excess Ile, Phe or Met each decreased TORC1 activity, and in the case of excess Met this response requires the Tco89 subunit. Met-triggered growth arrest required the kinase Gcn2, as well as its positive regulators the Gcn1-Gcn20 complex and its effector the Gcn4 transcription factor. Although Gcn2 is known to be activated by uncharged tRNAs, we show here that addition of excess Met triggers robust activation of Gcn2 and induction of Gcn4 expression, a response which is required to dampen TORC1 activity. Unexpectedly, we find that this response requires the endocytosis of the methionine transporter of Mup1. Taken together, our findings suggest that endocytic trafficking of amino acid transporters in response to excess amino acids initiates a response that involves activation of Gcn2 kinase followed by dampening of TORC1 activity, induction of autophagy, and G1 arrest. These findings suggest a potential mechanism for coupling endocytosis of an amino acid transporter to stress signaling for cell growth control.

## Results

### Specific amino acids trigger a slow growth program in yeast

While much is known about how yeast cells respond to nutrient deprivation less is known about how cells respond and adapt to excess amino acid availability. To explore how cells manage excess amino acid availability, we characterized yeast growth in the presence of defined minimal media (SCD) following addition of excess amino acids. In standard minimal media, wildtype *S. cerevisiae* cultures exhibit a lag phase, followed by exponential growth and a plateau associated with diauxic shift. This characteristic pattern was also observed when most amino acids are supplied in excess. However, certain amino acids, when supplied in excess, were severely inhibitory for growth, including Ile, Phe, Met and His (**FIG 1A**). The ability of His to inhibit growth has already been reported and is linked to copper homeostasis (Watanabe et al., 2014). We therefore decided to focus our investigation on growth inhibition by excess Ile, Phe, and Met.

**Figure 1.**
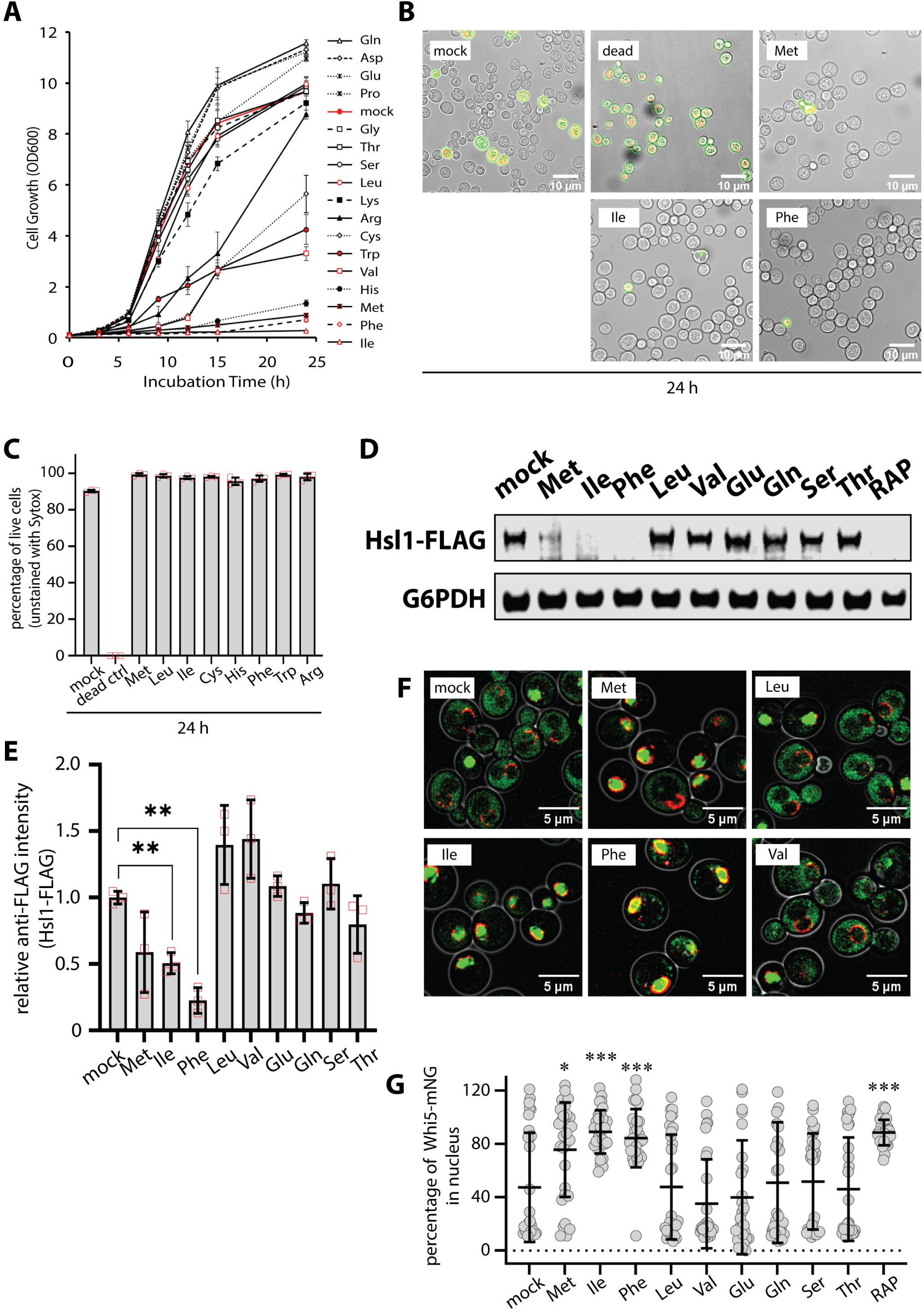
Specific amino acids trigger a slow growth program in yeast. (A) Yeast cells (strain background: SEY6210) were cultured overnight to mid-log phase (OD_600_ = 0.5) and then sub-cultured in fresh SCD media to initiate the experiment at OD_600_ = 0.05. Cells were grown over a 24 hour period in a shaking incubator at 30°C and OD_600_ was measured in a spectrophotometer at the indicated time points. (B) After culturing cells for 24 hours in the same conditions as (A), cells were stained with SYTOX and imaged by fluorescence microscopy and DIC overlay. (C) For each sample in (B), SYTOX-positive and SYTOX-negative cells were counted (n≥100). (D) Yeast cells expressing FLAG-tagged Hsl1 (chromosomally tagged) were cultured and stimulated with the indicated amino acids as described in (A). Yeast cell lysates were analyzed by SDS-PAGE and immunoblotting was performed for using α-FLAG antibodies and α-G6PDH antibodies as a loading control. (E) Quantification of immunoblotting results in (D) over three biological replicate experiments (n=3). (F) Yeast cells expressing Whi5-GFP (green) and Nup49-mCherry (red, nuclear envelope) were analyzed by fluorescence microscopy. (G) Quantification of results in (F). Total and nuclear Whi5-GFP signal were measured and a percentage nuclear signal was calculated for at least 30 cells in each condition tested (n≥30).

We first considered that addition of excess Ile, Phe, and Met may be toxic, resulting in loss of cell viability. To test this, we analyzed cultures using SYTOX viability dye, which stains dead cells. Yeast cells cultures in defined minimal media (SCD) for 24 hours exhibited a high degree of viability (9.7% SYTOX positive cells) while no viability was detected in cells heated at 65°C for 10 minutes (100% SYTOX positive cells). In contrast, cells grown in the presence of excess Ile, Phe, or Met exhibited increased viability (2.4%, 2.9%, and 0.7% SYTOX positive cells, respectively) compared to regular SCD (**FIG 1B-C**). These results indicate that excess Phe, Ile, or Met does not decrease yeast cell viability.

We next considered the possibility the presence of excess Ile, Phe and Met induces cell cycle arrest. To examine cell cycle arrest, we imaged the yeast septin ring by imaging cells expressing mCherry-tagged Shs1, a component of the spetin ring. After culturing cells for 24 hours in minimal media, most cells exhibited septin ring assemblies (**FIG S1A-B**). In contrast, addition of excess Ile, Phe, or Met resulted in a significant decrease in septin ring assemblies (**FIG S1A-B**) consistent with cell cycle arrest. Notably, immunoblot analysis of FLAG-tagged Shs1 revealed a doublet in minimal media that collapsed to a single species following addition of excess Ile, Phe or Met (**FIG S1C**) suggesting either decreased abundance or altered post-translational modifications. Proper septin ring assembly is required for transition to mitosis, and one important checkpoint is the kinase Hsl1 which localizes to septin rings and promotes the G2/M transition (Finnigan et al., 2016). Immunoblot analysis of lysates from yeast cells expressing FLAG-tagged Hsl1 revealed that excess Ile, Phe or Met resulted in decreased abundance of Hsl1-FLAG, while treatment with other amino acids either increased or did not significantly alter Hsl1-FLAG abundance (**FIG 1D-E**). We also examined subcellular localization of Whi5-GFP which localizes to the nucleus in early G1 but exits the nucleus at the onset of the G1/S transition. We observed a significant increase in cells exhibiting nuclear-localized Whi5-GFP following treatment with excess Ile, Phe, or Met (**FIG 1F-G**). Based on this analysis, we conclude that treatment of yeast cells with excess Ile, Phe or Met triggers G1 arrest.

### Activation of autophagy by amino acid surplus

In yeast, starvation is known to trigger G1 arrest as well as autophagy (An et al., 2014; Matsui et al., 2013). To test if excess amino acid stimulation triggers autophagy, we used a bulk autophagy reporter assay (Noda and Klionsky, 2008) and found that the same amino acids that slowed cell growth (Ile, Phe and Met) also induced bulk autophagy (**FIG 2A**). This induction of autophagy by Ile, Phe, and Met was not observed in *Δatg5* mutant cells, indicating this response is *bona fide* autophagy. The induction of autophagy by addition of excess amino acids was unexpected, since autophagy is usually associated with nutrient restriction rather than nutrient excess.

**Figure 2.**
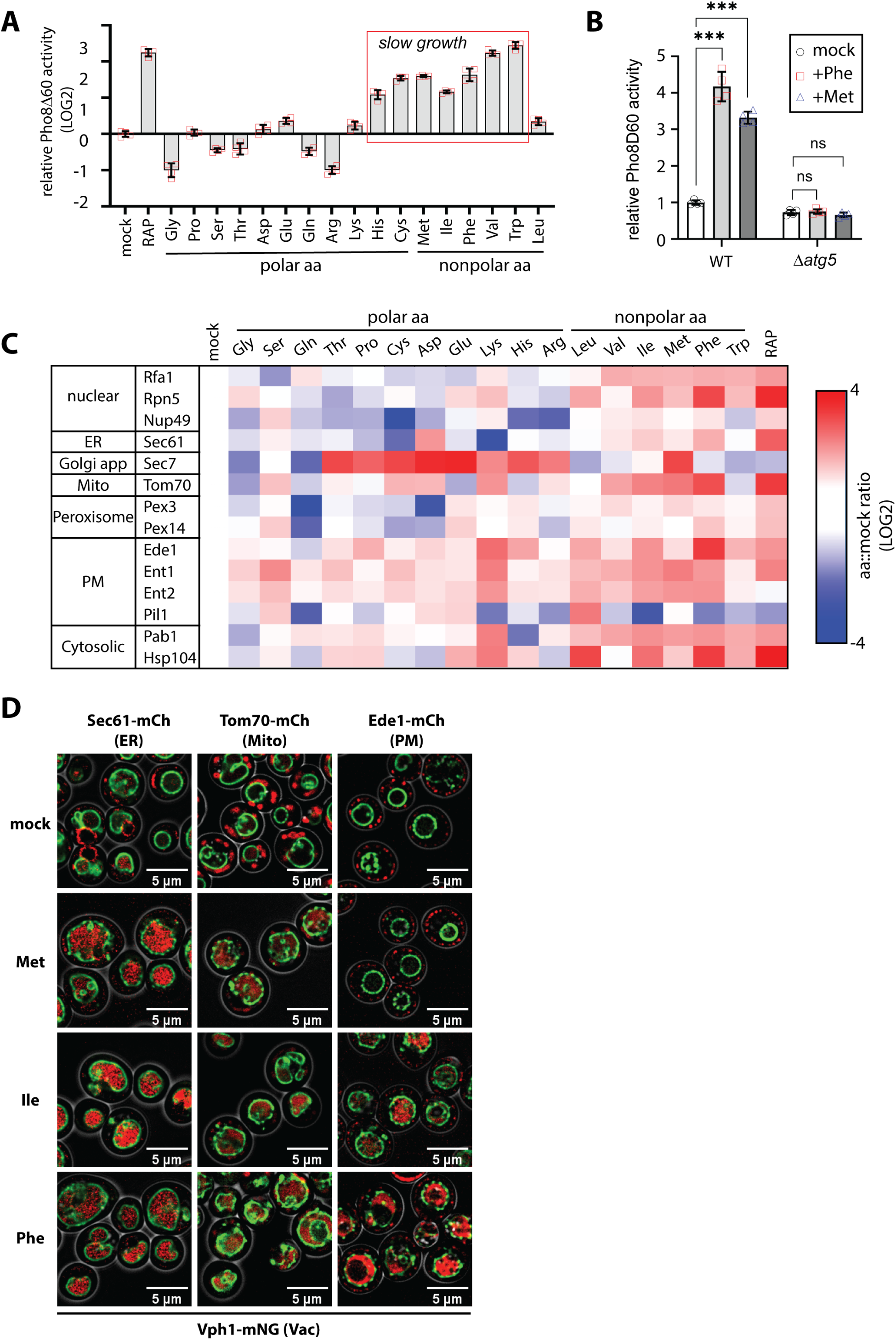
Activation of autophagy by amino acid supplementation. (A) Bulk autophagy was measured in yeast cells (strain background: SEY6210) using the Pho8Δ60 activity reporter assay with the indicated conditions. Assays were preformed in biological triplicate expeirments (n=3) for each indicated condition. (B) Bulk autophagy assays were performed as in (A) comparing wildtype (WT) and *Δatg5* mutant cells. (C) Yeast cells expressing Vph1-mNG (vacuole membrane) along with the indicated mCherry-tagged organelle marker (left column) were analyzed for fraction vacuole localization in individual cells (n≥30 cells per condition). The color indicated in each position of the heat map is the average fraction vacuole localization, normalized to control (mock) condition. Red indicates an increase in vacuole localization compared to control conditions, whereas blue indicates a decrease in vacuole localization compared to control conditions. (D) Fluorescence microscopy images of data representative of the heat map shown in (C).

To further explore this finding, we systematically characterized cargo selective autophagy in response to different amino acids when supplied in excess. Specifically, we used fluorescence microscopy to quantify the localization of different autophagy cargo to the lumen of the vacuole. This analysis is displayed as a heat map (**FIG 2C**) where the vacuole localization of each cargo in different conditions was quantified over a population of cells (n≥30) and normalized to the mock condition (i.e., regular minimal media). As a positive control, cells were treated with rapamycin (RAP) which resulted in broad induction of autophagy (**FIG 2C**). Notably, treatment of cells with excess Ile or Phe triggered autophagy of various cargo with a pattern resembling RAP (**FIG 2C-D** and **FIG S2**). Met stimulation also triggered autophagy, albeit with a slightly different pattern (**FIG 2C-D**).

To our knowledge, mitophagy is not known to be triggered by excess amino acids. To better understand this response, we performed kinetic analysis of Tom70 localization to the vacuole lumen which differences between the different amino acid stimuli. For example, excess Phe triggered a sight but significant induction in mitophagy as early as 3 hours following stimulation, and the magnitude of this effect became greater over a 24 hour stimulation time course (**FIG 3A**). In contrast, excess Met triggered a delayed response that was significant only after 12 hours of stimulation (**FIG 3B**) and exhibited titratable activation of mitophagy up to 1.2mg/mL (**FIG S3A**). To validate these results we measured vacuolar localization of other mitochondrial markers including Om45 (outer mitochondrial membrane), Cox4 (inner mitochondrial membrane), and Mdh1 (matrix). In each case the presence of excess Phe, Met, and to a lesser extent Ile triggered localization to the vacuole lumen (**FIG 3C**). Confirming that Ile, Phe and Met trigger mitophagy via the conventional pathway, we found that Tom70 localization to the vacuole lumen triggered by excess Ile, Met, and Phe is dependent on ATG5, which is required for autophagosome formation, as well as ATG11 and ATG32 (**FIG 3D-E**), which are part a mitophagy adaptor complex (Schuster and Okamoto, 2022). Notably, SILAC-based quantitative proteomic analysis revealed that stimulation with excess methionine increased association between Atg11 (bait) and other autophagy factors such as Atg1, Atg17, Atg29 and Atg31 (**FIG 3F**). Taken together, these findings indicate that excess Phe, Ile and Met can serve as triggers to activate mitophagy in yeast cells.

**Figure 3.**
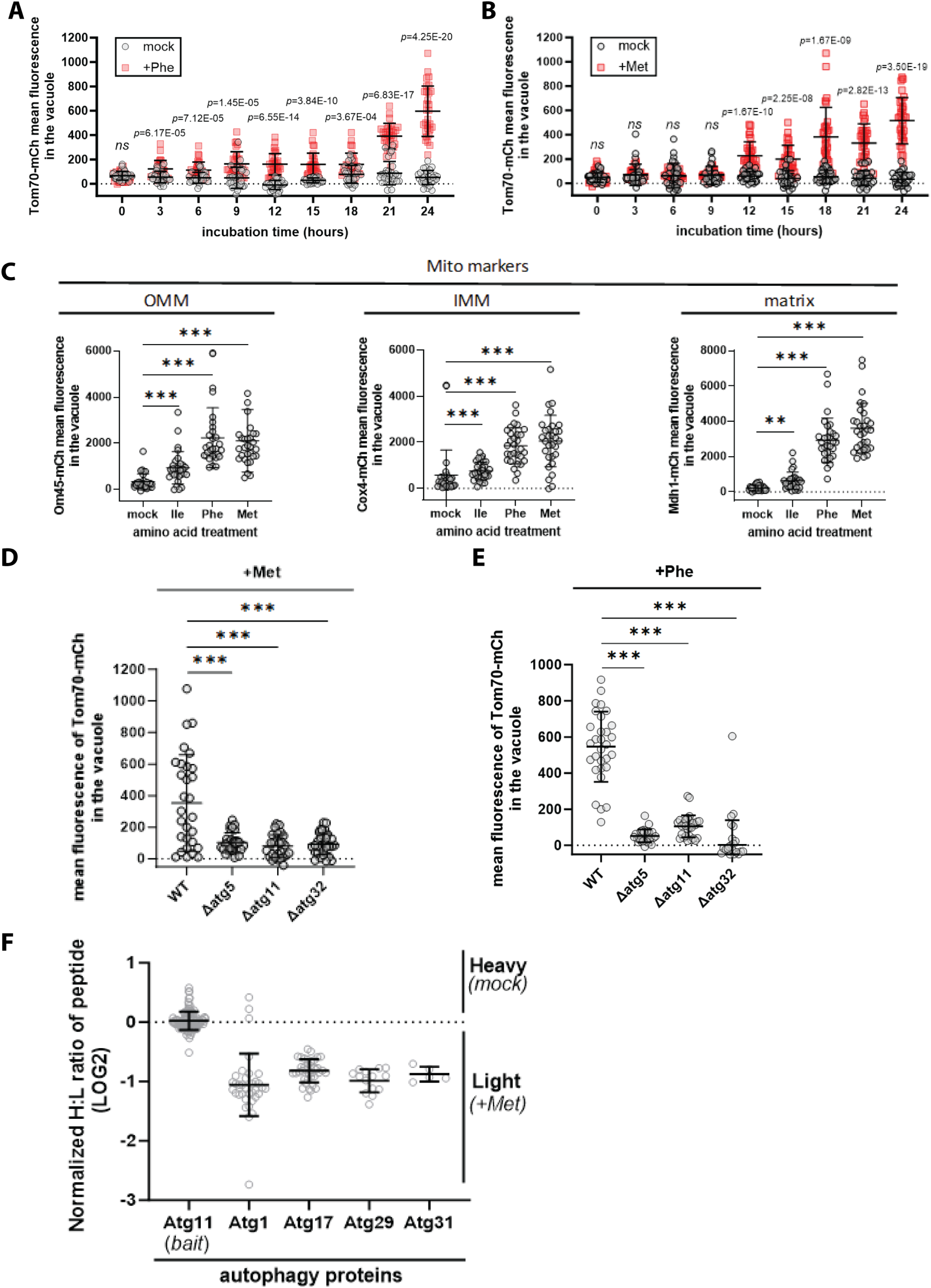
Activation of mitophagy by amino acid supplementation. (A) For individual cells expressing Vph1-mNG (vacuole membrane) and Tom70-mCherry (mitochondria), mean fluorescence in the vacuole was measured for the indicated time points and conditions (n≥30 cells). (B) For individual cells expressing Vph1-mNG (vacuole membrane) and Tom70-mCherry (mitochondria), mean fluorescence in the vacuole was measured for the indicated time points and conditions (n≥30 cells). (C) For individual cells expressing Vph1-mNG (vacuole membrane) and the indicated mitochondrial markers mean fluorescence in the vacuole was measured for the indicated conditions (n≥30 cells). (D) For individual cells expressing Vph1-mNG (vacuole membrane) and Tom70-mCherry (mitochondria), mean fluorescence in the vacuole was measured for the indicated conditions (n≥30 cells). (E) For individual cells expressing Vph1-mNG (vacuole membrane) and Tom70-mCherry (mitochondria), mean fluorescence in the vacuole was measured for the indicated conditions (n≥30 cells). (F) SILAC-MS analysis was performed on affinity-purified Atg11-FLAG (bait) comparing cells grown in standard media (SCD) and cells grown following methionine supplementation.

### Regulation of TORC1 signaling by excess methionine

The observed cellular response to excess Ile, Phe, and Met includes a G1 arrest and induced autophagy, which is similar to a starvation response. Indeed, the cargo-selective autophagy pattern induced by excess Phe and Ile mimicked that observed by treatment with rapamycin (**FIG 2C**) suggesting that these amino acids may interfere with TORC1 signaling. To test this hypothesis, we examined TORC1 activity in minimal defined media (SCD) following stimulation with excess amino acids. As expected, treatment with rapamycin inhibited TORC1 activity, while providing all amino acids in excess increased TORC1 activity (**FIG 4A-B**). While supplying excess of most amino acids had no detectable affect on TORC1 signaling, excess Ile, Phe and Met inhibited TORC1 activity (**FIG 4A-B**). This effect was most significant with excess Ile and more partial with excess Phe and Met (**FIG 4A-B**). Time course analysis following stimulation with excess Met revealed partial inhibition of TORC1 activity within 3 hours which could be eliminated altogether by co-treatment with rapamycin (**FIG 4C-D**) indicating that excess Met dampens but does not completely inhibit TORC1 activity.

**Figure 4.**
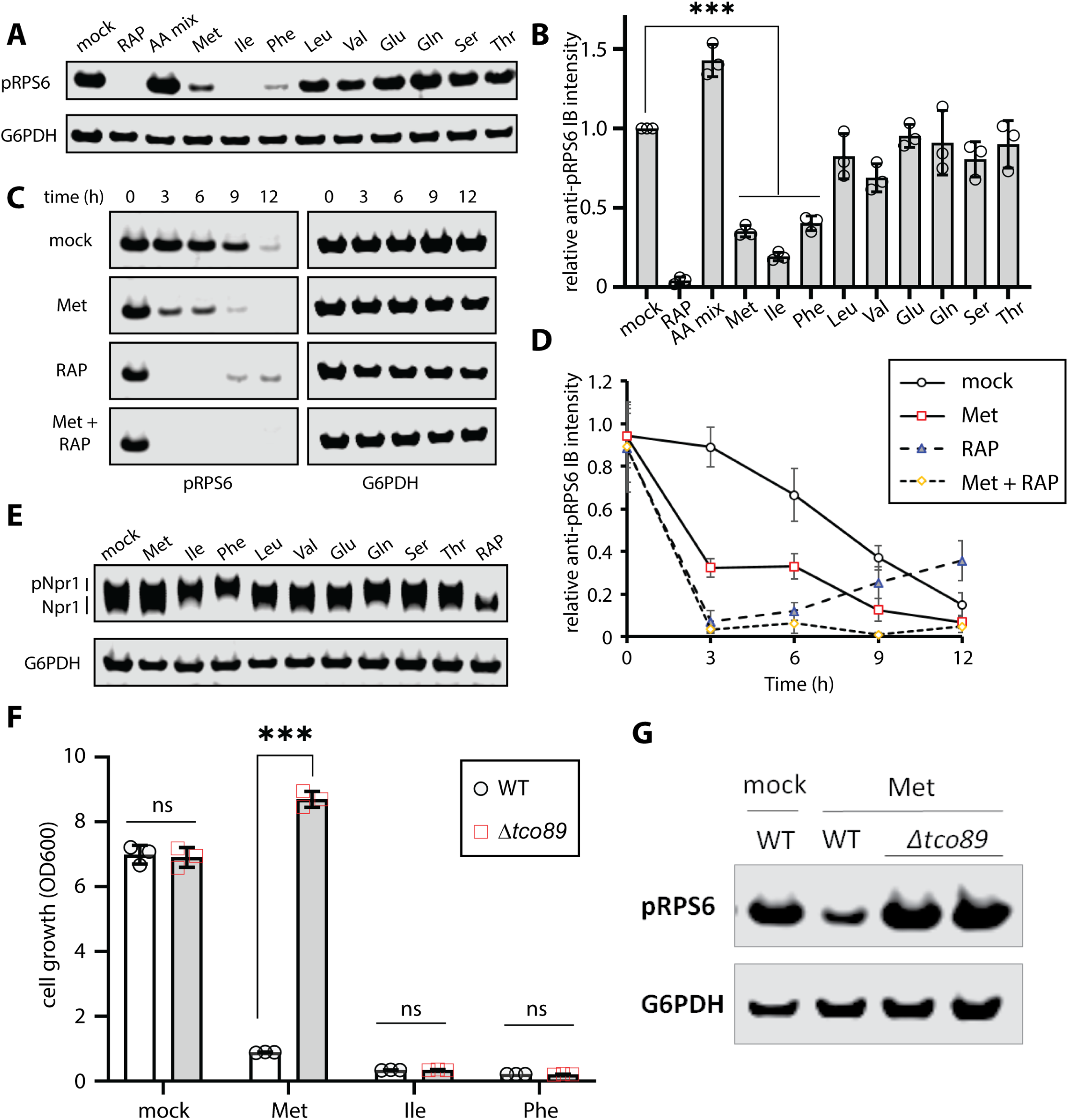
Amino acid supplementation dampens TORC1 activity. (A) Cells cultured in standard media (SCD) or with the indicated treatment were collected and cell lysates were analyzed by SDS-PAGE and immunoblotting for pRPS6 as a readout for TORC1 activity. G6PDH was used as a loading control. (B) Quantification of the immunoblotting results shown in (A) over three biological replicate experiments (n=3). (C) Cells cultured in standard media (SCD) or with the indicated treatment were collected and cell lysates were analyzed by SDS-PAGE and immunoblotting for pRPS6 as a readout for TORC1 activity. G6PDH was used as a loading control. (D) Quantification of the immunoblotting results shown in (C) over three biological replicate experiments (n=3). (E) Cells cultured in standard media (SCD) or with the indicated treatment were collected and cell lysates were analyzed by SDS-PAGE and immunoblotting for Npr1-HA as a readout for TORC1 activity. G6PDH was used as a loading control. (F) Analysis of yeast overnight growth with the indicated culture conditions (n≥3). (G) Cells cultured in standard media (SCD) or with the indicated treatment were collected and cell lysates were analyzed by SDS-PAGE and immunoblotting for pRPS6 as a readout for TORC1 activity. G6PDH was used as a loading control.

Unexpectedly, we noted that Tco89, a non-essential subunit of the TORC1 kinase complex, is required for Met-triggered growth arrest, but not Ile- or Phe-triggered growth arrest (**FIG 4F**). Consistent with this result, Tco89 is required for Met-triggered mitophagy but not for mitophagy triggered by rapamycin (**FIG S4A-D**). Tco89 is also required for Met-triggered dampening of TORC1 kinase activity (**FIG 4G**). These results indicate that Tco89 is a negative regulator of TORC1 activity in the context of excess methionine.

### Methionine activates Gcn2 and the ISR to dampen TORC1 signaling

To better understand what factors mediate the response to methionine, we screened for yeast knockout strains that fail to mediate growth arrest in the presence of excess Met. Notably, mutations disrupting autophagy did not prevent growth arrest in the presence of excess Met (**FIG 5A**), indicating that autophagy is a feature of this response but not a requirement to arrest growth. Interestingly, we found that loss of Gcn2, the kinase that phosphorylates eIF2α during starvation, as well as its upstream regulators Gcn1 and Gcn20 prevented Met-triggered growth arrest (**FIG 5A**). Gcn4, a downstream effector of the Gcn2 response that mediates the transcriptional response to starvation, is also required for Met-triggered growth arrest (**FIG 5A**). These results suggest that the Gcn2 pathway senses and responds to the presence of excess methionine. Consistent with this finding, this pathway is also required for Met-triggered mitophagy (**FIG S5A-B**). To test if the Gcn2 pathway is required for Met-triggered dampening of TORC1 activity, we examined the TORC1 response to excess Met in mutants defective for the Gcn2 response. In each of these mutants (*Δgcn1*, *Δgcn20*, *Δgcn2, Δgcn4*) excess Met failed to dampen TORC1 activity (**FIG 5B-C**). This result indicates that the Gcn2 pathway is required for TORC1 dampening.

**Figure 5.**
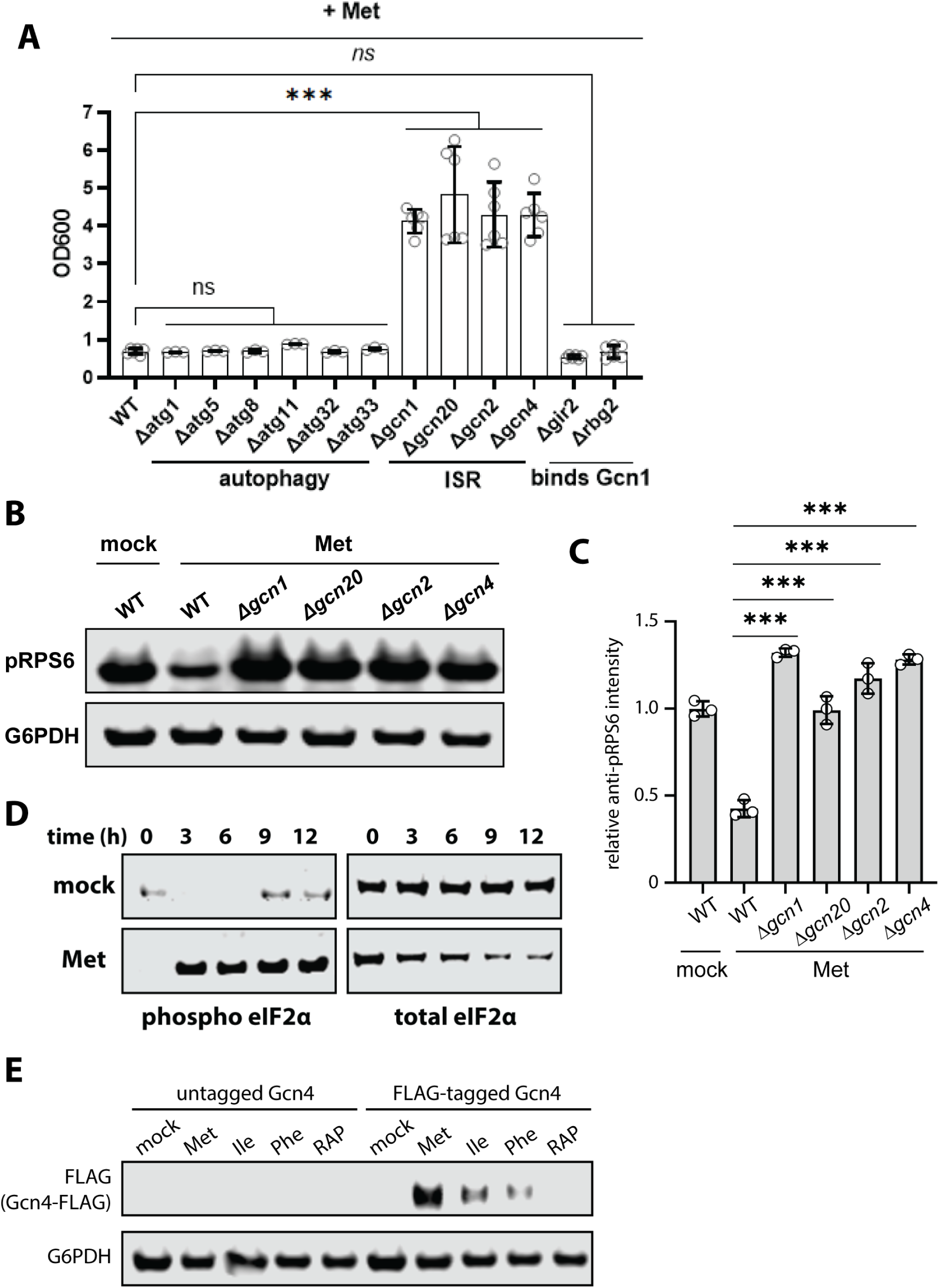
Methionine activates Gcn2 to slow growth. (A) Analysis of yeast overnight growth with the indicated culture conditions (n≥3). (B) Cells cultured in standard media (SCD) or with the indicated treatment were collected and cell lysates were analyzed by SDS-PAGE and immunoblotting for pRPS6 as a readout for TORC1 activity. G6PDH was used as a loading control. (C) Quantification of the immunoblotting results shown in (B) over three biological replicate experiments (n=3). (D) Cells cultured in standard media (SCD) or with the indicated treatment were collected and cell lysates were analyzed by SDS-PAGE and immunoblotting for phospho-eIF2α as a readout for Gcn2 activity. Total eIF2α was used as a loading control. (E) Cells harboring chromosomal FLAG-tagged Gcn4 were cultured in standard media (SCD) or with the indicated treatment were collected and cell lysates were analyzed by SDS-PAGE and immunoblotting for Gcn4-FLAG expression as a readout for Gcn2 activity. G6PDH was used as a loading control.

To test if Met stimulates Gcn2 activity, we performed immunoblot analysis of eIF2α, which can be phosphorylated at Ser51 by Gcn2 to dampen global levels of translation initiation. Addition of excess Met stimulated robust eIF2α phosphorylation at Ser51 within 3 hours (**FIG 5D**). Consistent with this result, we also observed induced expression of Gcn4 within one hour of Met stimulation (**FIG 5E**). Combined, these results indicate that excess Met promotes signaling through the Gcn2 pathway.

### Endocytosis is required for Met-triggered activation of Gcn2

The Gcn2/Gcn4 pathway is activated by amino acid deprivation and by ribosome collisions (Misra et al., 2024) so it was unexpected that this pathway is also activated by stimulation with excess Met. If Gcn2 is responding to increased availability of Met in the cytosol, we reasoned that activation should correlate with Met transporter activity. Several studies have shown that the high affinity Met transporter Mup1 is stably expressed at the plasma membrane in media lacking Met, but undergoes rapid endocytosis and vacuolar trafficking following addition of Met to the media (Hepowit et al., 2023a; Hepowit et al., 2023b; Lee et al., 2019; Lin et al., 2008). We hypothesized that blocking this endocytic response and stabilizing Mup1 at the plasma membrane would facilitate increased Met transport and hyper-sensitize cells to Met-triggered growth arrest. To test this, we examined yeast strains expressing Mup1 fused to UL36, a deubiquitylase enzyme that prevents ubiquitin modification and therefore endocytic trafficking (Hepowit et al., 2023a; MacDonald et al., 2012; Stringer and Piper, 2011). Unexpectedly, fusion of Mup1 to catalytically active UL36, but not a catalytic dead variant, prevented Met-triggered growth arrest (**FIG 6A**). By comparison, fusing the UL36 to Hse1 (a subunit of ESCRT-0), which prevents cargo sorting by the ESCRT pathway resulting in cargo accumulation on endosomal membranes (Lee et al., 2017; Stringer and Piper, 2011), did not prevent Met-triggered growth arrest (**FIG 6A**). This result suggests that endocytosis of Mup1, but not ESCRT-mediated delivery to the vacuole lumen, is required to mediate Met-triggered growth arrest. Importantly, cells expressing Mup1-UL36 failed to induce Gcn4 following exposure to methionine (**FIG 6C-D**) consistent with the hypothesis that Mup1 endocytosis activates Gcn2. However, we cannot exclude the possibility that the Mup1-UL36 fusion protein may have trans effects on other cargo or protein complexes at the plasma membrane.

**Figure 6.**
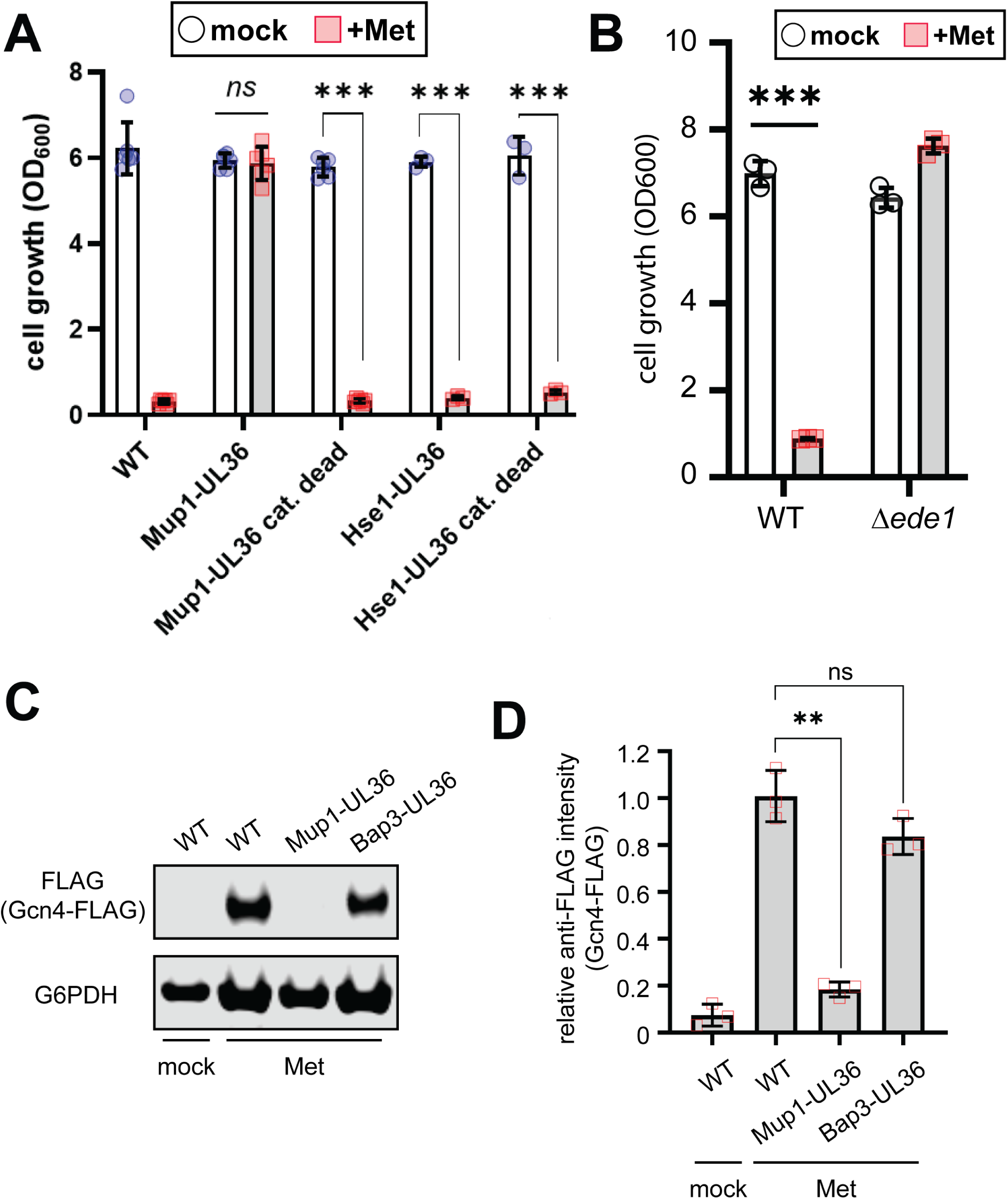
Endocytosis is required for Met-triggered Gcn2 activation. (A) Analysis of yeast overnight growth with the indicated culture conditions (n≥3). (B) Analysis of yeast overnight growth with the indicated culture conditions (n≥3). (C) Cells harboring chromosomal FLAG-tagged Gcn4 were cultured in standard media (SCD) or with the indicated treatment were collected and cell lysates were analyzed by SDS-PAGE and immunoblotting for Gcn4-FLAG expression as a readout for Gcn2 activity. G6PDH was used as a loading control. (D) Quantification of immunoblotting results shown in (C) from three biological replicate experiments (n=3).

Based on these observations, we hypothesized that endocytosis of Mup1, rather than influx of Met into the cytosol, contributes to activation of Gcn2. Consistent with this hypothesis, the endocytic adaptor Ede1, which is required for Met-triggered Mup1 endocytosis (Hepowit et al., 2023b; Wrasman et al., 2018), is required for Met-induced growth arrest (**FIG 6B**). Similarly, Ppz phosphatases, which are also required for Met-triggered Mup1 endocytosis (Lee et al., 2019), are required are required for Met-induced growth arrest (**FIG S6A**), TORC1 dampening (**FIG S6B**), and Gcn4 induction (**FIG S6C-D**). These results provide additional evidence that endocytosis plays a role in the activation of Gcn2 in response to Met.

## Discussion

During nutrient deprivation, cells respond by arresting the cell cycle and shifting to activities that conserve resources such as autophagy. In this study, we report that stimulation with excess amino acids – specifically Ile, Phe, and Met – triggers a response that mimics starvation. This is characterized by activation of the Gcn2 kinase, dampening of TORC1 activity, induction of autophagy and G2 arrest. Thus, amino acid surplus triggers a response that in many ways parallels a starvation response. Other published studies have reported evidence that similar effects may occur in mammalian cells. In K562 cells, high concentrations of phenylalanine were reported to inhibit growth in a manner that is dependent on the LAT transporter (Sanayama et al., 2014). In liver cancer cells, methionine supplementation was reported to trigger slow growth (Tripodi et al., 2020). Notably, cancer-derived cell lines are frequently addicted to methionine, a phenomenon known as the Hoffman effect (Kaiser, 2020), indicating that physiological responses to Met availability are altered in cancer.

Our study also characterizes the relationship between amino acid surplus and cargo-specific autophagy. As noted, Phe, Ile and Met were activators of bulk autophagy and this was validated through analysis of several cargo-specific autophagy pathways. Phe and Ile triggered cargo-specific autophagy patterns that mimicked rapamycin treatment, which is consistent with the observed TORC1 inhibition by those amino acids. Met also induced autophagy, but with a slightly different pattern. The cargo-selective autophagy heatmap (**FIG 2C**) revealed other notable patterns. For example, certain polar amino acids induced Golgi-phagy. It is unclear why these induced Golgi-phagy, but one speculative possibility is that surplus of certain amino acids might somehow damage specific membranes in the cell. Another interesting observation is that basic amino acids such as His and Arg generally inhibited autophagy. This is consistent with one study that reported excess Arg inhibits autophagy in breast cancer cells (Sannino et al., 2023) suggesting that at least some of the patterns on this heat map are likely to be conserved in mammalian cells.

Gcn2 activation is most often associated with starvation, but it is also activated by other stresses including UV irradiation and oxidative stress (Misra et al., 2024). Activation of Gcn2 by starvation conditions is proposed to occur via binding of uncharged tRNAs to a domain related to the histidyl tRNA synthetase, whereas activation of Gcn2 can also occur during translation stress via interactions with collided ribosomes (Misra et al., 2024). Additional studies will be needed to determine if either of these mechanisms contribute to the methionine-triggered Gcn2 activation we report in this study, but we hypothesize that endocytosis of amino acid transporters may signal to activate Gcn2 in certain conditions. This may involve regulation of upstream factors, such as the Gcn1-Gcn20 complex. Future studies will need to address the relationship between methionine levels, methionine transporters, and the mechanism by which these lead to regulation of the Gcn2 pathway.

## Materials and methods

### Yeast strains and growth conditions

All yeast strains used in this study were haploid cells derived from *Saccharomyces cerevisiae* SEY6210 (listed in Supplemental Table S1) using standard genetic and molecular biology techniques. Epitope tags and fluorescence proteins were fused at the C-terminus of target proteins by chromosomal integration. Gene deletions were generated by swapping target ORFs with antibiotic resistance marker gene through homologous recombination. Strains with combined gene knockout and epitope-fusion genotypes were generated by mating and sporulation, isolated by tetrad dissection, and confirmed by diagnostic PCR. Unless otherwise stated, yeast cells were grown in minimal SCD (detailed recipe in Supplemental Table S2) at 30ᵒC with agitation at 220 rpm. Precultures were obtained from freshly streaked YPD (yeast-peptone-dextrose) plate, grown for 16 h, subcultured into fresh media at starting OD600 of 0.1, and allowed to grow to early log phase (OD600 of 0.3 - 0.4) prior to treatment. Cultures in minimal SCD were treated with surplus amino acid (2 mg/ml amino acid supplementation) or rapamycin (0.2 µg/ml). Cells for amino acid starvation were precultured to mid-log (OD600 of 0.6 – 0.8), harvested by centrifugation, washed twice with sterile dH2O, and inoculated into culture medium containing YNB (6.7 g/L, without amino acids) and 2% glucose. Cell growth was determined by measuring the optical density (OD600) of cells after 24 h of incubation from an initial culture of 0.1 OD600.

### Fluorescence microscopy

Yeast cells endogenously expressing fluorescent fusion proteins (mCherry, mNeonGreen, BFP) were concentrated by centrifugation (3,500 X *g* for 30 s) and directly visualized using a DeltaVision Elite Imaging system [Olympus IX-71 inverted microscope; Olympus 100× oil objective (1.4 NA); DV Elite sCMOS camera, GE Healthcare] equipped with softWoRx software (v7.0.0; GE Healthcare). Images obtained from the red (Alexa Fluor 594; 475 nm excitation, 523 nm emission) and green (FITC; 575 nm excitation, 632 nm emission) filter channels were deconvolved using softWoRx, and the mean fluorescence intensity units in the vacuole (expressed as integrated density fluorescence over the area of the vacuole) were measured using Fiji (Schindelin et al., 2012).

### Protein preparation and immunodetection

Proteins from 5.0 OD600 units of cells were precipitated with 10% trichloroacetate in Tris-EDTA buffer (10 mM Tris-HCl [pH8.0], 1 mM EDTA) in LoBind microfuge tube (Eppendorf, Hamburg, Germany), washed with 100% acetone, resuspended in 100% acetone by sonication, lyophilized by vacuum centrifugation, solubilized in 100 µl cracking buffer (50 mM Tris [pH 7.5], 1 mM EDTA, 1% SDS, 6 M urea) and 100 µl 2X urea sample buffer (150mM Tris [pH 6.8], 6 M urea, 6% SDS, 40% glycerol, bromophenol blue), and resolved by electrophoresis on a NuPAGE (4-12% or 8% Bis-Tris) gel. Proteins were transferred onto PVDF membranes (0.45 µm, GE Healthcare Amersham) by electrophoretic transblotting, blocked with 3% bovine serum albumin in TBST (20 mM Tris-HCl [pH 7.4], 150 mM NaCl, 0.05% Tween-20), and incubated with the following primary antibodies: mouse anti-FLAG (1:1000; Sigma-Aldrich; clone M2; F3165; RRID AB_262044), mouse anti-HA (1:1000; ThermoFisher Scientific, 12CA5 monoclonal antibody), rabbit anti-eIF2α (Cell Signaling Technology), rabbit anti-phospho-Ser51 eIF2α (Cell Signaling Technology), rabbit phospho-Ser235/236 S6 ribosomal protein (Cell Signaling Technology), and rabbit anti-G6PDH (1:10,000; Sigma-Aldrich; A9521; RRID AB_258454). The secondary antibodies used were IRDye 680RD-goat anti-rabbit (LI-COR) and IRDye 800CW-goat anti-mouse (LI-COR). Fluorescence of blots was visualized using the Odyssey CLx Imaging System (LI-COR) and quantified using Image Studio Lite (LI-COR).

### Nonspecific autophagy assay

All strains used for nonspecific bulk autophagy assay endogenously express the Pho8Δ60 alkaline phosphatase (ALP) variant. To generate these strains, the genomic *PHO8* gene was replaced with the *PHO8*Δ*60*::*NATMX* DNA cassette by homologous integration at the *PHO8* locus to loop out the native gene. After autophagy induction by 12 h treatment with 2 mg/ml amino acid supplement or 0.2 µg/ml rapamycin, cells at 4 OD600 units were harvested by centrifugation (200 X *g* for 3 min), resuspended with 0.2 ml ice-cold ALP assay buffer (250 mM Tris-HCl [pH 9.0], 10 mM MgSO4, 10 µM ZnSO4), and lysed with 0.2 ml glass beads by vortex agitation (1 min vortex, 1 min rest [6 cycles]). Lysates were diluted by addition of 0.2 ml ice-cold ALP assay buffer and clarified by centrifugation (21,100 X *g* for 1 min). Protein concentrations were determined by bicinchoninic acid assay (BCA, Pierce BCA Protein Assay Kit). Reaction mixture (50 µl lysate, 450 µl ALP assay buffer, 50 µl ALP substrate [55 mM α-naphthyl phosphate in ALP assay buffer]) was incubated at 30ᵒC for 20 min and stopped with 0.5 ml 2 M glycine-NaOH (pH 11.0). Fluorescence was measured using a 96-well plate reader at 345 nm excitation and 472 nm emission.

### Mass spectrometry

Protein interactome of Atg11 was determined by LC-MS/MS as previously described (Hepowit et el., 2020) with minor modification. Briefly, *Δarg4Δlys2* yeast cells endogenously expressing Atg11-FLAG were grown to early log phase (OD600 of ∼0.4) in SILAC SCD with either light or heavy Arg and Lys, and treated with 2 mg/ml Met or Phe for 12 h. Cells were harvested by centrifugation (3,500 X *g* for 10 min) and disrupted by bead beating using ice-cold lysis buffer (50 mM HEPES [pH 7.4], 150 mM NaCl, 5 mM EDTA, 0.1% NP-40, 10 mM iodoacetamide, 1 mM 1,10-phenanthroline, 1X EDTA-free protease inhibitor cocktail [Roche], 1 mM phenylmethylsulfonyl fluoride, 20 µM bortezomib, 1X PhosStop [Roche], 10 mM NaF, 20 mM BGP, and 2 mM Na_3_VO_4_). Lysate was clarified by centrifugation at 21,100 × *g* for 10 min at 4°C and supernatant was transferred into a new tube with 50 µL of EZview anti-FLAG M2 resin slurry (Sigma) and incubated for 2 hr at 4°C with rotation. The resin was washed three times with cold wash buffer (50 mM HEPES [pH 7.4], 150 mM NaCl, 0.05% NP-40) and resin-binding proteins were eluted with 90 µL elution buffer (100 mM Tris-HCl, pH 8.0, 1% SDS) by heating 98°C for 5 min. The collected eluate was reduced with 10 mM DTT, alkylated with 20 mM iodoacetamide, and precipitated with 300 µL PPT solution (50% acetone, 49.9% ethanol, and 0.1% acetic acid). Light and heavy protein pellets were dissolved with Urea-Tris solution (8 M urea, 50 mM Tris-HCl, pH 8.0). Heavy and light samples were combined, diluted four-fold with water, and digested with 1 µg MS-grade trypsin (Gold, Promega) by overnight incubation at 37°C. Peptides were purified using a SEPAK C18 column (washed with 0.1% acetic acid, and recovered in 200 µl of 80% acetonitrile 0.1% acetic acid), lyophilized by vacuum centrifugation, and resolubilized in 12 µl 0.1% trifluoroacetic acid. A 0.5 µl peptide solution was injected into a capillary reverse-phase analytical column (360 μm O.D. × 100 μm I.D.) using a Dionex Ultimate 3000 nanoLC and autosampler and analyzed using a Q Exactive Plus mass spectrometer (Thermo Scientific). Data collected were searched using MaxQuant (ver. 1.6.5.0).

## Acknowledgements

We are grateful to A Ebert and T Martinez for technical advice and feedback. NLH and JAM were supported by NIH grant R35 GM144112 (to JAM).

## Supplementary Figure Legends

**Figure S1.**
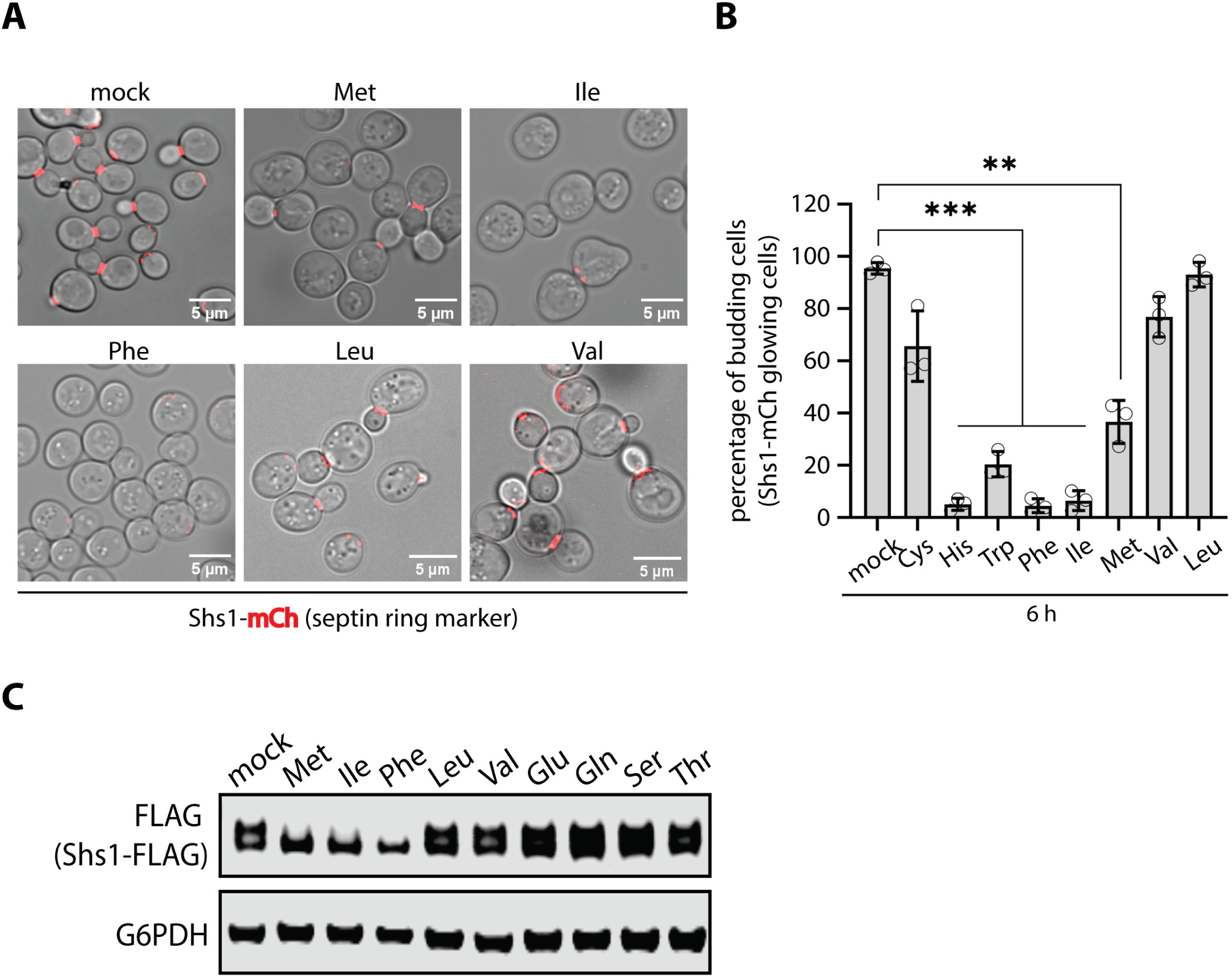
Specific amino acids inhibit septin assembly. (A) Yeast cells expressing Shs1-mCherry (red, septin ring) were analyzed by fluorescence microscopy and overlayed with DIC images. (B) Quantification of the results shown in (A) over three biological replicate experiments (n=3). (C) Yeast cells expressing Shs1-FLAG (chromosomally tagged) were cultured and stimulated with the indicated amino acids. Cell lysates were analyzed by SDS-PAGE and immunoblotting was performed for using α-FLAG antibodies and α-G6PDH antibodies as a loading control.

**Figure S2.**
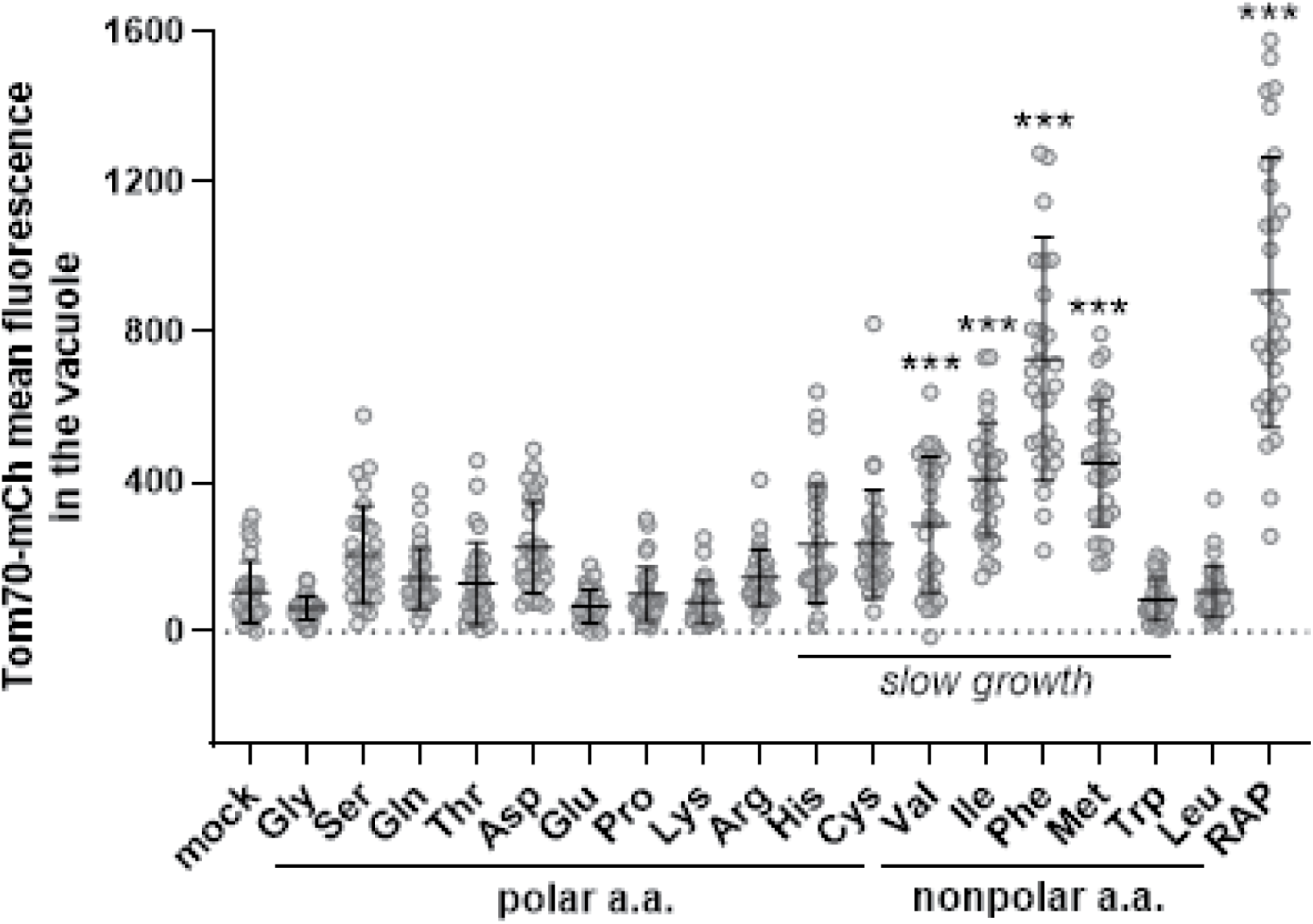
Activation of mitophagy by amino acid supplementation. For individual cells expressing Vph1-mNG (vacuole membrane) and Tom70-mCherry (mitochondria), mean fluorescence in the vacuole was measured for the indicated time points and conditions (n≥30 cells).

**Figure S3.**
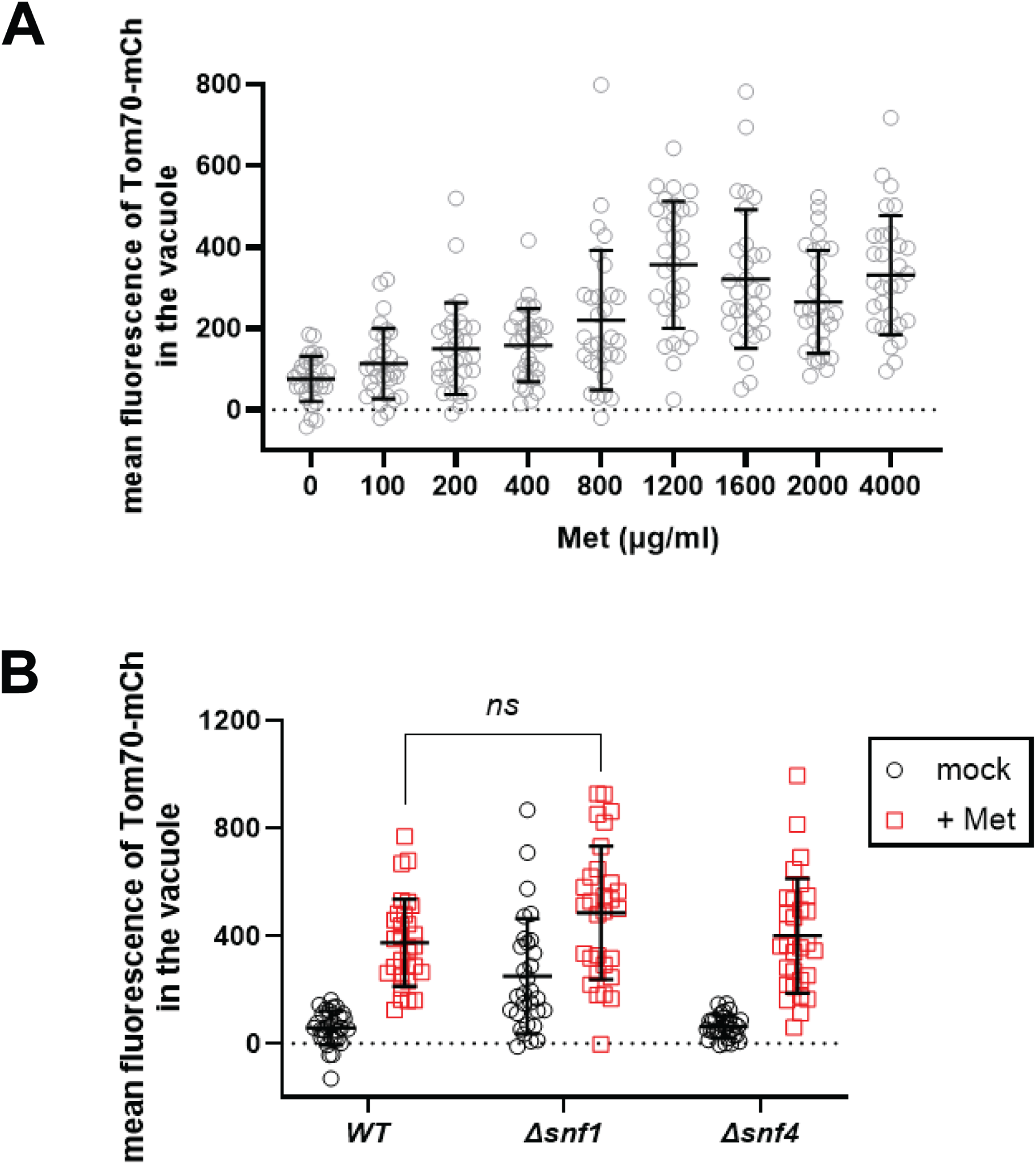
Features of Met-triggered mitophagy. (A) For individual cells expressing Vph1-mNG (vacuole membrane) and Tom70-mCherry (mitochondria), mean fluorescence in the vacuole was measured for the indicated methionine concentrations (n≥30 cells per condition). (B) For individual cells expressing Vph1-mNG (vacuole membrane) and Tom70-mCherry (mitochondria), mean fluorescence in the vacuole was measured for the indicated time points and conditions (n≥30 cells).

**Figure S4.**
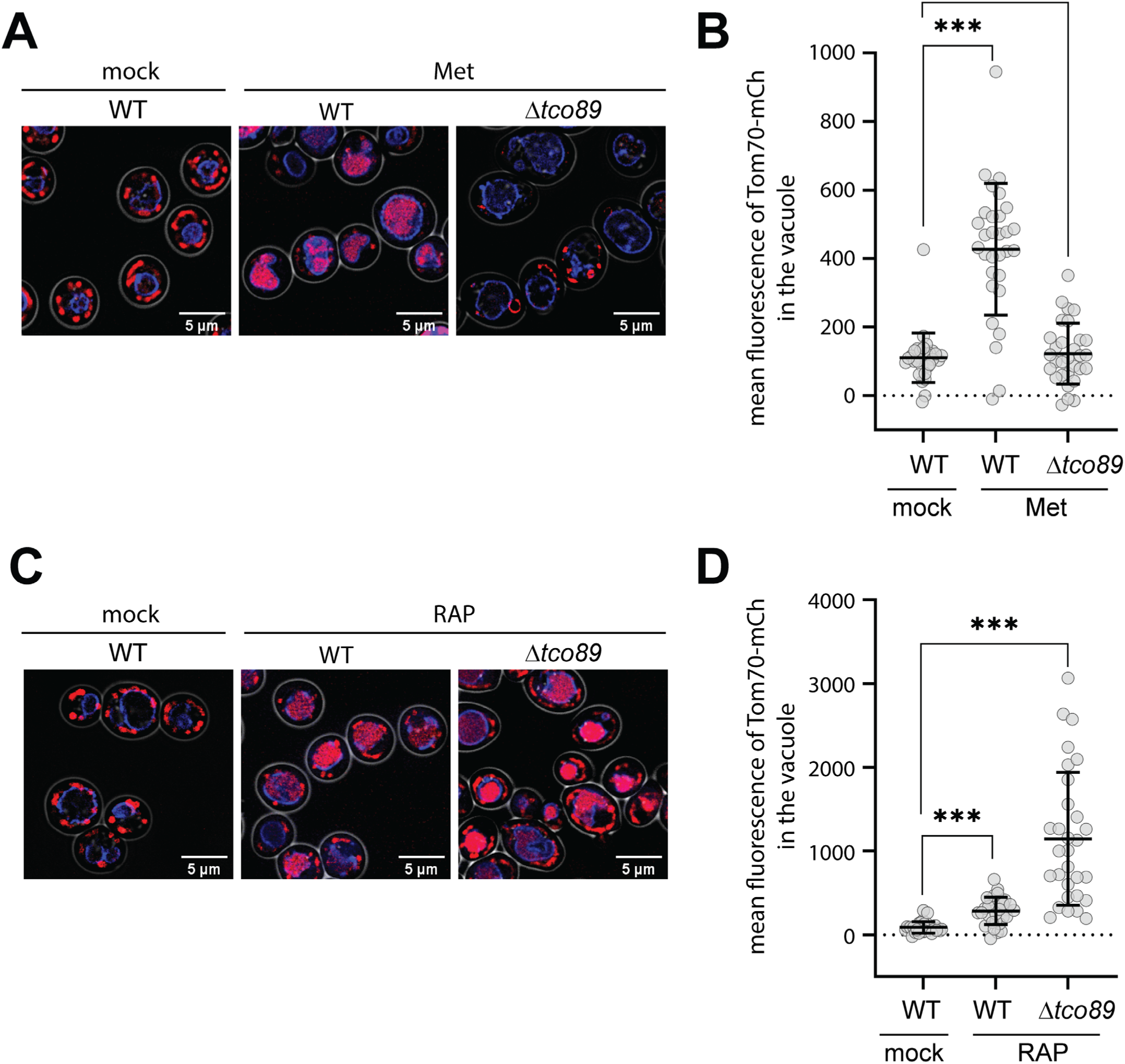
Tco89-dependent activation of mitophagy by Met supplementation. (A) Cells expressing Vph1-mTagBFP2 (vacuole membrane, blue) and Tom70-mCherry (mitochondria, red) were imaged by fluorescence microscopy and DIC (overlayed). (B) For individual cells from the experiment shown in (A), mean Tom70-mCherry fluorescence in the vacuole was measured for the indicated time points and conditions (n≥30 cells). (C) Cells expressing Vph1-mTagBFP2 (vacuole membrane, blue) and Tom70-mCherry (mitochondria, red) were imaged by fluorescence microscopy and DIC (overlayed). (D) For individual cells from the experiment shown in (A), mean Tom70-mCherry fluorescence in the vacuole was measured for the indicated time points and conditions (n≥30 cells).

**Figure S5.**
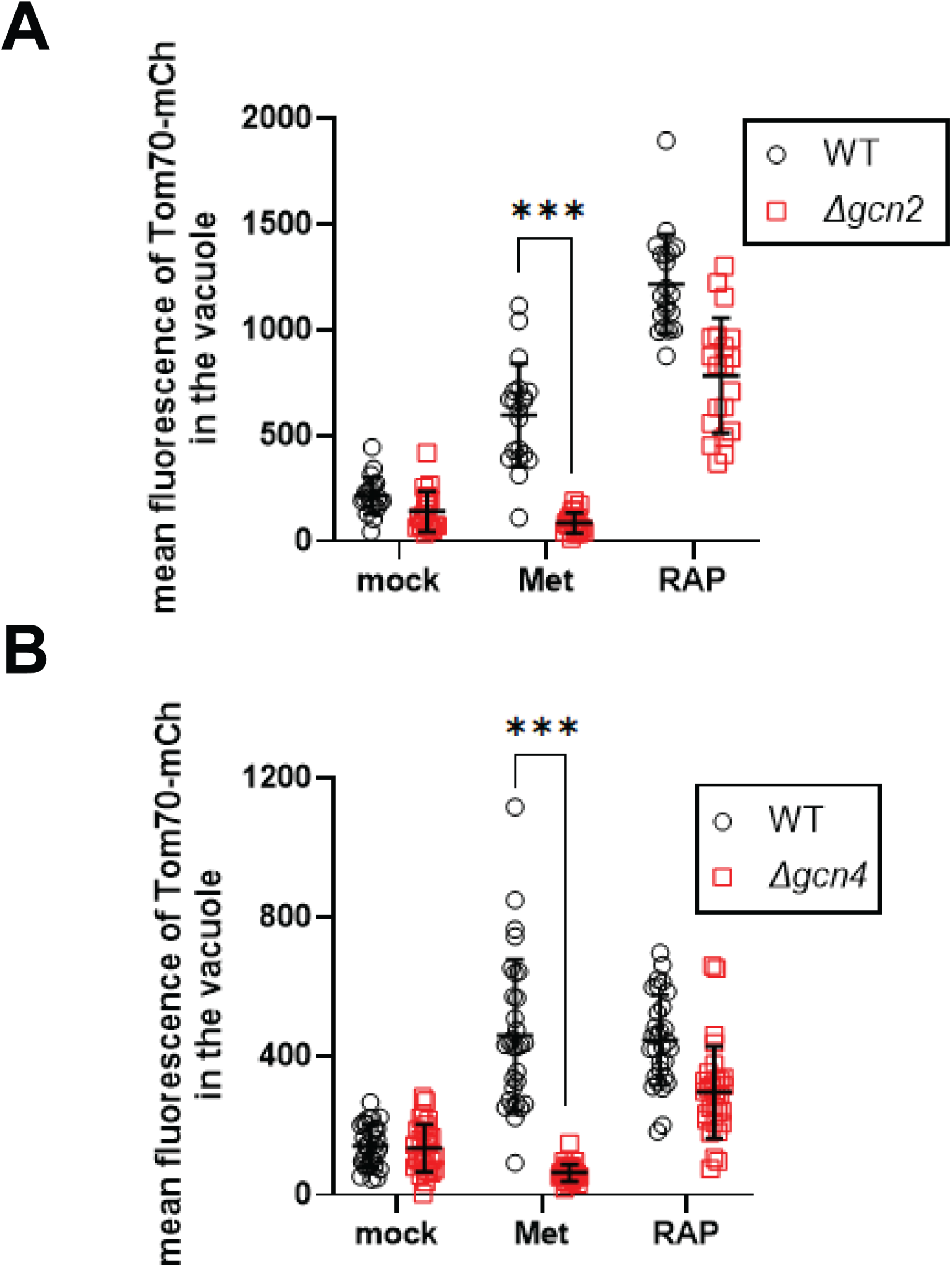
Met-triggered mitophagy is dependent on the Gcn2 pathway. (A) For individual cells expressing Vph1-mNG (vacuole membrane) and Tom70-mCherry (mitochondria), mean fluorescence in the vacuole was measured for the indicated time points and conditions (n≥30 cells). (B) For individual cells expressing Vph1-mNG (vacuole membrane) and Tom70-mCherry (mitochondria), mean fluorescence in the vacuole was measured for the indicated time points and conditions (n≥30 cells).

**Figure S6.**
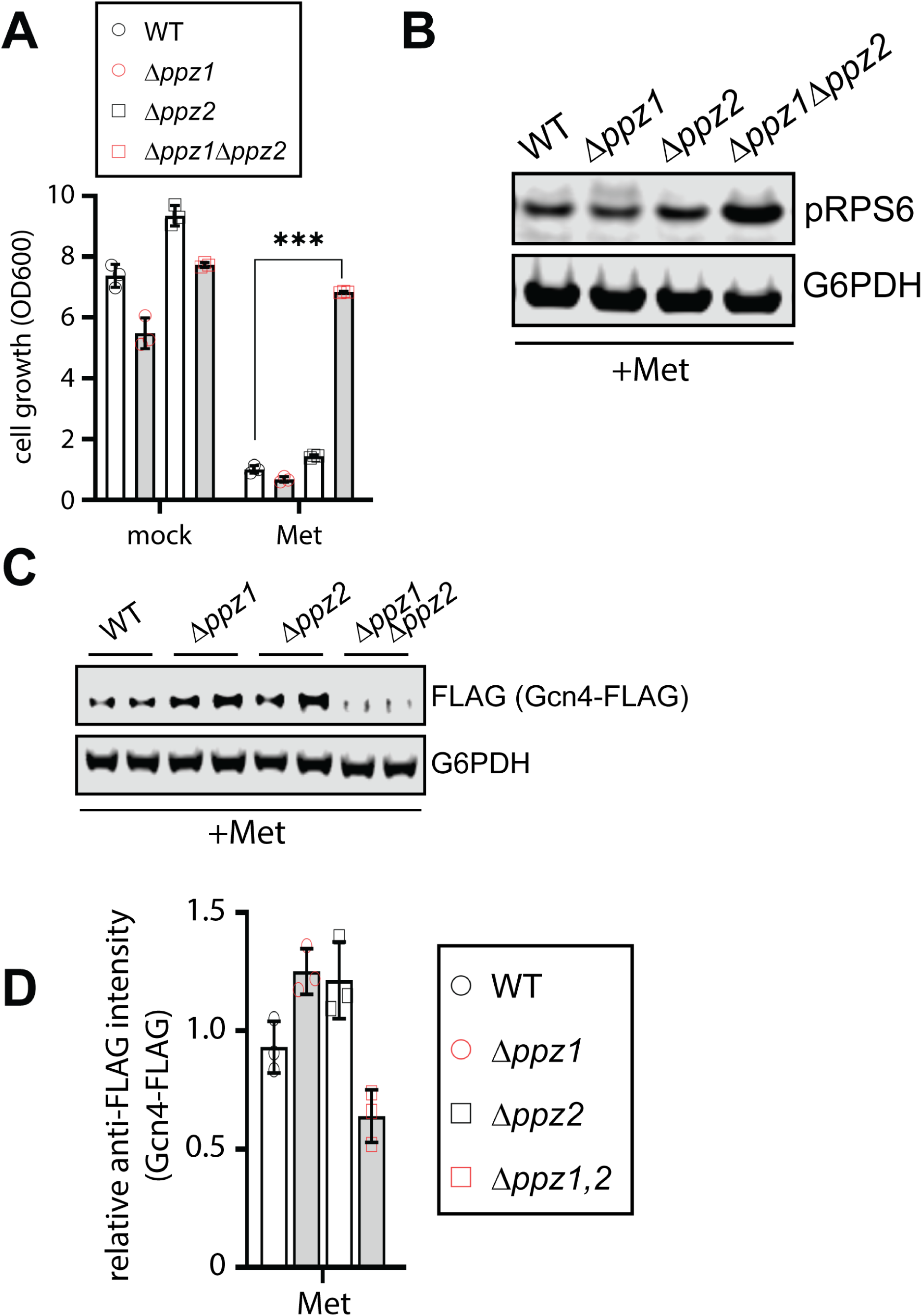
Ppz phosphatases are required for Met-triggered Gcn2 activation. (A) Analysis of yeast overnight growth with the indicated culture conditions (n≥3). (B) Cells cultured in standard media (SCD) or with the indicated treatment were collected and cell lysates were analyzed by SDS-PAGE and immunoblotting for pRPS6 as a readout for TORC1 activity. G6PDH was used as a loading control. (C) Cells harboring chromosomal FLAG-tagged Gcn4 were cultured in standard media (SCD) or with the indicated treatment were collected and cell lysates were analyzed by SDS-PAGE and immunoblotting for Gcn4-FLAG expression as a readout for Gcn2 activity. G6PDH was used as a loading control. (D) Quantification of immunoblotting results shown in (C) from three biological replicate experiments (n=3).

